# A cooperatively breeding mouse shows flexible use of its vocal repertoire according to social context

**DOI:** 10.1101/2024.05.07.592648

**Authors:** Léo Perrier, Aude de Witasse-Thézy, Aurélie Pradeau, Carsten Schradin, Michael D. Greenfield, Nicolas Mathevon, Florence Levréro

**Affiliations:** ENES Bioacoustics Research Laboratory, CRNL, CNRS, Inserm, University of Saint-Etienne, Saint-Etienne, France; Université de Strasbourg, CNRS, IPHC UMR 7178, F-67000 Strasbourg, France; School of Animal, Plant & Environmental Sciences, University of the Witwatersrand, Private Bag 3, WITS 2050, Johannesburg, South Africa; Department of Ecology and Evolutionary Biology, University of Kansas, Lawrence, KS 66045 USA; Institut universitaire de France; Ecole Pratique des Hautes Etudes, CHArt Lab, PSL University, France

**Keywords:** Ultrasonic vocalizations, social communication, vocal repertoire, turn-taking

## Abstract

Mice exchange information using chemical, visual and acoustic signals. Long ignored, mouse ultrasonic communication is now considered to be an important aspect of their social life, transferring information such as individual identity or stress levels. However, whether and how mice modulate their acoustic communications is largely unknown. Here we show that a wild mouse species with a complex social system controls its vocal production both qualitatively and quantitatively, depending on social context. We found that the African striped mouse *Rhabdomys pumilio*, a cooperatively breeding species, has a vocal repertoire consisting of seven call types, which it uses differently depending on whether the individuals encounter another mouse that is familiar, unfamiliar, of the same or different sex. Familiar individuals, whether of the same or different sex, vocalize more than two unfamiliar same-sex individuals. The greatest diversity of vocalisations is recorded when a female and a male first encounter, suggesting that certain calls are reserved for courtship. Our results highlight that familiar mice alternate their vocalisations (turn-taking) while unfamiliar individuals tend to overlap one another. These observations suggest that African striped mice control the production and temporal dynamics of their vocalisations, addressing targeted information to specific receivers via the acoustic channel.

## Introduction

In recent decades, our knowledge about mammal vocal communication has exploded, and now enables us to understand how information can be exchanged via sound between individuals in a number of contexts, such as vocal recognition vocal recognition (1,2), mate choice (3,4), antagonist interactions (5,6), and so on (7). However, studies have mainly focused on a small number of mammal groups, with primates and marine mammals probably leading the way (8,9). Rodents, which account for over 40% of all the mammal species on Earth (10), have thus received belated attention (11–16). Although we know that rodents, most of which are highly social, vocalize to exchange information between individuals in many contexts, such as aggression, play, exploration, etc. (11,17,18), our knowledge therefore remains rudimentary. It has to be said that studying their ultrasonic communications requires highly specific equipment and know-how that is relatively difficult to master. Yet understanding the characteristics of acoustic communication in rodents is essential if we are to grasp the foundations of their social life.

A particularly interesting rodent subfamily is that of the Murinae, the old World mice and rats, due to their major biomedical significance (19,20). The species that have begun to be studied, however, remain are few in number, with most studies focusing concentrating on laboratory strains of house mice and of brown rats (*Mus musculus* and *Rattus norvegicus*). Recording contexts are generally poorly representative of the diversity of social contexts encountered by free-living wild animals (21). In fact, rodent vocalisations are still widely regarded as mere markers of an individual’s emotional state (22) and it is not known to what extent mice can flexibly control the emission of their ultrasounic vocalisations.

Describing and understanding the biological significance of the vocal repertoire of rodents requires associating the acoustic characteristics of vocalisations with the contexts in which they are produced. The classic approach, which has been widely used to describe the vocal repertoires of laboratory rats and mice, is to form categories of vocalisations, based on frequency and temporal criteria that are simple to evaluate. Rat vocalisations are usually classified into two categories corresponding to two different frequency ranges, around 22 kHz (calls made in aversive contexts) and around 50 kHz (calls made in positive contexts; 11–13). These two call categories can be divided into several subcategories depending on the context of emission and on the level of arousal (26,27). Laboratory mice also produce a variety of calls, which have been classified according to their acoustic parameters, in particular their frequency shape (28). Unlike rats, it is difficult to attribute clear emotional characteristics to the ultrasonic vocalisations of laboratory mice. However, researchers have recently studied variations in call types as a function of overall social context, and demonstrated that females produce longer, more complex calls than males (29). These variations in the proportion of different call type emitted may differ between mouse strains rather than between social contexts (30). In wild house mice, studies have shown that males produce more USVs in the presence of female urine than male urine, and even more so if the female is unfamiliar (31). With regard to interactions with animals of the same sex, males are less vocal in response to other male presence’s cue, and females are more likely to communicate with a familiar female compared to an unfamiliar one (31–33). While these experiments report only quantitative changes in USVs, researchers have recently begun to focus on the repertoire of wild house mice, and show that some variations in frequency modulations can be induced by presenting different individuals behind a Plexiglass cover (34). These vocalisations also play a role in female attraction (35–37), but do not seem to induce oestrus (38). In rodents outside the family Murinae, e.g. voles or singing mice, vocalisations also mediate social interactions (39,40).

Cataloguing and classifying the vocalisations that define the repertoire of a rodent species often remains a challenge. Unlike the discrete vocal repertoire of many bird species, composed of different types of vocalisations that are relatively easy to distinguish acoustically (e.g. zebra finch; 29), the vocal repertoire of a mouse or rat is graded: the acoustic boundaries between vocalisations categories are blurred and have been subject to the more or less subjective appreciation of the observer (42). To define vocalisation categories, authors often simply use thresholds based on a few acoustic parameters (e.g. average call duration or frequency). Recently, advances in artificial intelligence have made it possible to describe a vocal repertoire more objectively. Deep learning approaches now support the exploration of large recording banks in an unsupervised manner (43–45). In the present study, we used such machine learning techniques dedicated to describing the vocal repertoire of the house mouse and rat (29,46,47) to understand complexity of the ultrasound repertoire and its use in diverse social contexts in a wild rodent.

We studied acoustic communication in a wild strain of the African striped mouse *Rhabdomys pumilio*, a murine rodent like the laboratory mouse and rat. This mouse species has been studied extensively in the wild and has a complex social system (48,49). The African striped mouse is a key study model for understanding the links between rapid variations in environmental constraints and their consequences on social life: this species demonstrates social flexibility, with the ability to alternate between solitary life and life in family groups, depending on the level of reproductive competition and population density (50). Groups typically consist of one breeding male, 2-4 breeding females and up to 25 adult offspring of both sexes (51). Social structure in family groups predicts numerous and diverse interactions between all group members (51,52), in other words a vocal communication network rich in information exchange. Moreover, the striped mouse is diurnal and easily observed both in captivity and in the field (51,53). Thus, African striped mice offer a valuable opportunity to explore acoustic communication in a species phylogenetically close to the laboratory mouse, but being entirely wild and displaying a richer and more flexible social life.

Here we tested the hypothesis that the vocalisations produced by the African striped mouse differ according to social context, depending on the sex and familiarity of the interacting individuals. We conducted an unsupervised machine learning classification of the vocalisations produced in different social contexts. We also measured the precise timing of acoustic communication to evaluate occurrence vs avoidance of call overlap when two mice vocalized, an index that could reflect a ’turn-taking’ propensity in these rodents. Our results suggest that striped mice control their vocal production both qualitatively and quantitatively.

## Methods

### Ethics

All procedures were approved by the Loire Ethic Committee for Animal Experimentation of the University of Saint-Etienne (CEEAL-UJM n°98) under reference #35444-2022021412408695.

### Subjects

We performed 45 pairwise encounters with a total of 87 African striped mice (3 mice were used in two different encounters). All animals were born in captivity and descended from individuals captured in 2016 in the Goegap Nature Reserve in South Africa. In our laboratory, the animals are housed in pairs (breeding pair) or unisex groups (siblings) in Zolux Neo Panas XL cages (71*41*48 cm). Enrichments (tubes, wooden houses and platforms) are regularly installed, moved and changed in their cage. The mice are kept under a 12/12h light-dark cycle (lights on at 08:00 to 20:00). The experiments were carried out between 10:00 am and 14:00 pm. Mice were fed 90 minutes before the start of the experiment with 4 g of seeds’ mix for rodents (Zolux©).

### Experimental design

The pairwise encounters were carried out with 2 adult individuals (aged from 4 to 8 months), either familiar or unfamiliar. For each of these categories, we carried out encounters between two males, two females or two individuals of different sex. Familiar encounters between males and females were carried out with pairs of individuals housed in the same cage for at least 3 weeks and with no dependent offspring. Familiar encounters between individuals of the same sex were carried out with brothers or sisters housed together since birth. Unfamiliar encounters (individuals of the same or different sex) were carried out with two individuals born and housed in two separate rooms, with no possibility of having seen, heard or smelled each other since birth.

For each pairwise trial in each part of the experiment, the two individuals were taken out of their maintenance cage and isolated separately in two other cages, which were placed in separated acoustic chambers for the hour preceding the experiment. The mice were then introduced into an experimental cage placed in another acoustic chamber (SilentBox© with acoustic foam-covered walls). The experimental cage comprised two compartments separated by an opaque, acoustically insulated partition, meaning animals were not able to hear each other before the encounter (one compartment per mouse; Zolux Neo Panas XL cage, 71*41*48 cm). The partition was effective, as no USVs were recorded before the encounter. A trapdoor allowed communication between the two compartments, which opening of the trapdoor was controlled from outside the acoustic chamber, avoiding any disturbance by human intervention. The cage floor was covered with 1 cm of clean bedding (Zolux Chambiose Nature). After a 20-minute habituation period, during which the mice remained in their respective compartments, the trapdoor was opened. Once one of the two mice had joined the other in its compartment, the two individuals remained together for 10 minutes, during which their vocal production was recorded (Figure 1).

**Figure 1.**
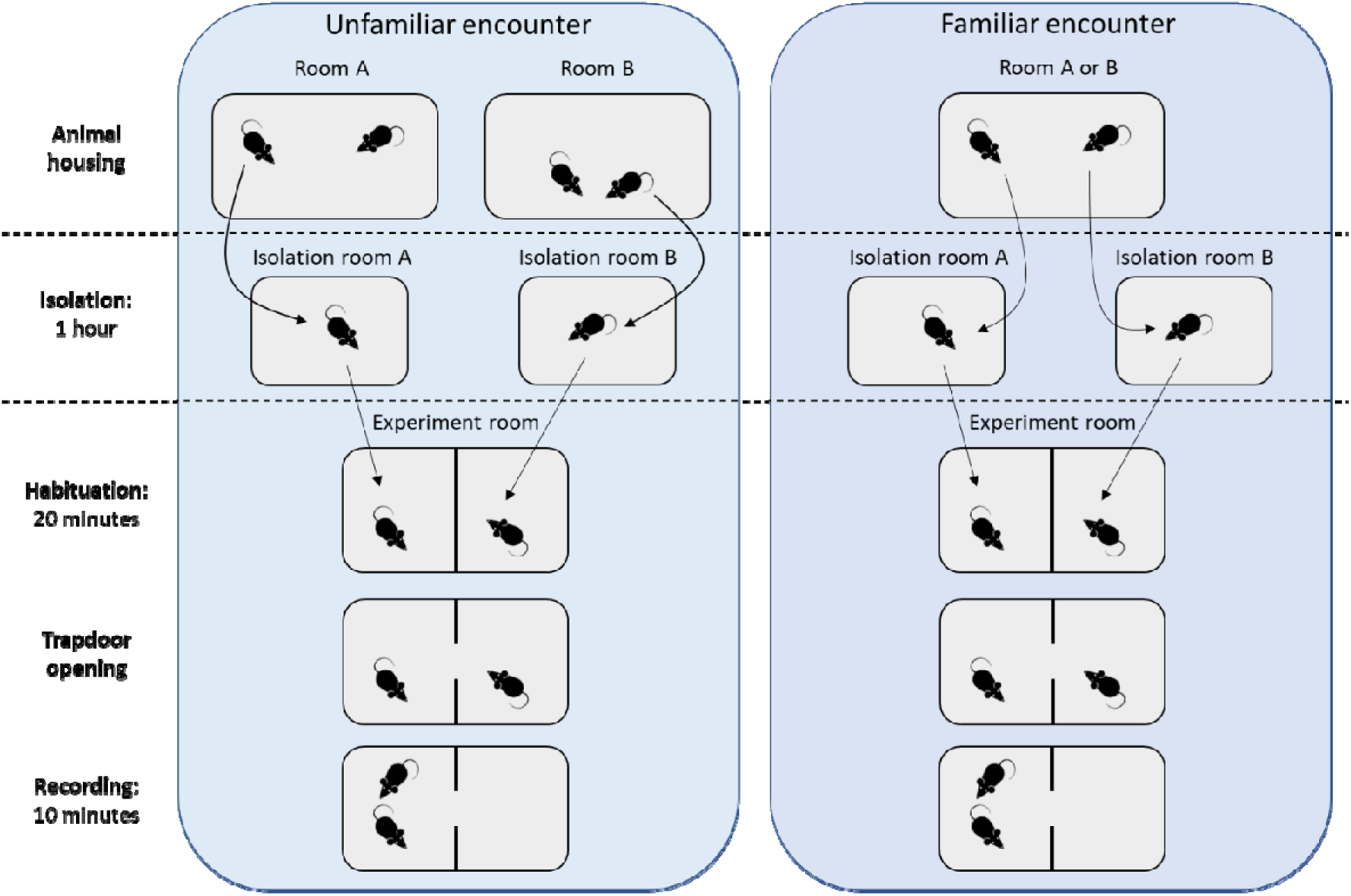
Encounter experiments. Left panel: encounter of unfamiliar individuals (housed in different rooms). Right panel: encounter of familiar individuals (housed in the same cage).

### Acquisition and detection of ultrasonic vocalisations

We recorded the ultrasonic vocalisations of the mice with an Avisoft© Bioacoustics CM16 microphone connected to a sound gate (Avisoft© Ultrasound Gate 416H, sampling rate = 300 kHz). The microphone was placed above the experimental cage. We performed the acoustic analysis using Deepsqueak© v3.0, a MATLAB software dedicated to the automatic detection and analysis of ultrasonic rodent vocalisations (43). Spectrograms were generated with a 0.0056-s window, 90% overlap and 0.0056 s NFFT. Two neural networks already trained in Deepsqueak© were used to detect vocalisations (Rat Detector YOLO R1 and Mouse Detector YOLO R2). To decide on the best parametrization to optimize the accuracy of automatic detection by the software, we first visually inspected the spectrograms of three randomly selected encounter experiments. Based on this test phase, we chose to set the tonality (Wiener entropy of the detected call) and the neural network score thresholds at 0.25 and 0.55 respectively.

### Classification of vocalisations

When vocalizing, mice generally emitted a series of vocalisations (usually called "calls") separated by silences. Using Deepsqueak© software, we isolated and analysed each vocalisation produced during each recording sequence (average: 159 vocalisations per minute of recording). Each vocalisation was described by a set of 9 acoustic descriptors representing its spectro-temporal characteristics (frequency contour, duration and average frequency of vocalisations; Table 1).

**Table 1.** Acoustic parameters used to describe the mice calls.

| Acoustic feature | Description |
| --- | --- |
| Call Length (s) | Duration of the call |
| Principle Frequency (kHz) | Median frequency of the call contour |
| Lowest Frequency (kHz) | Lowest frequency of the call contour |
| Highest Frequency (kHz) | Highest frequency of the call contour |
| Frequency Bandwidth (kHz) | Total frequency modulation of the call contour |
| Frequency Standard Deviation (kHz) | Standard deviation of the frequency of the call contour |
| Slope (kHz/s) | Mean slope of the call contour |
| Tonality | Wiener entropy of the call contour |
| Sinuosity | Length between the first and last points of the call contour divided by the Euclidean distance between the first and last points |

We then classified these vocalisations according to the 9 acoustic descriptors, using Deepsqueak’s unsupervised k-means clustering method. For this classification, we chose to double the weights given to the parameters describing the frequency contour (maximum amplitude for each time point in the spectrogram), compared with the weight for vocalisation duration and mean frequency. We adopted this method emphasizing spectral features because frequency modulation is expected to be particularly significant in mammalian vocal communication (54).

### Quantifying the temporal overlap between vocalisations

We defined calls as overlapping when the spectrogram presented two different frequencies at the same time. Because Deepsqueak detected overlapping calls as one single call, we first used the software’s unsupervised classification which assigned most of these paired calls to a novel cluster, different than non-overlapping calls. We then visually reviewed calls of this novel cluster to exclude non-overlapping ones that had been included due to classification errors.

### Visualization of data and statistics

To visualize mouse vocalisations in a two-dimensional acoustic space, we applied the UMAP algorithm (a non-linear algorithm of dimension reduction) on the 9 acoustic predictors used to classify vocalisations using “uwot” package (55)

For statistical analyses of the effect of social context on the number, temporal dynamics (number of vocalisations / minutes) and overlapping patterns of vocalisations, we used Bayesian multilevel models fitted with the “brms” R package with default priors (56). Posterior distributions of fitted values were discussed using their medians and 95% credible intervals (CIs).

## Results

### Vocal production depends on familiarity and sex

Figure 2 shows the average number of vocalisations recorded over 10 minutes for each type of encounter (66,137 vocalisations for all 45 trials). Mice made significantly fewer vocalisations during encounters between same-sex unfamiliar individuals compared to the other types of contexts (mean number of calls recorded over 10 minutes = 699 and 598, respectively for two unfamiliar females and two unfamiliar males’ encounters; comparison with other conditions: 95% CI [-2166; -442], 95% CI [- 2325; -603]). The number of vocalisations emitted was not significantly different between all other conditions, i.e. all encounters between familiar individuals, whether of the same sex or not, and encounters between unfamiliar individuals of the opposite sex.

**Figure 2.**
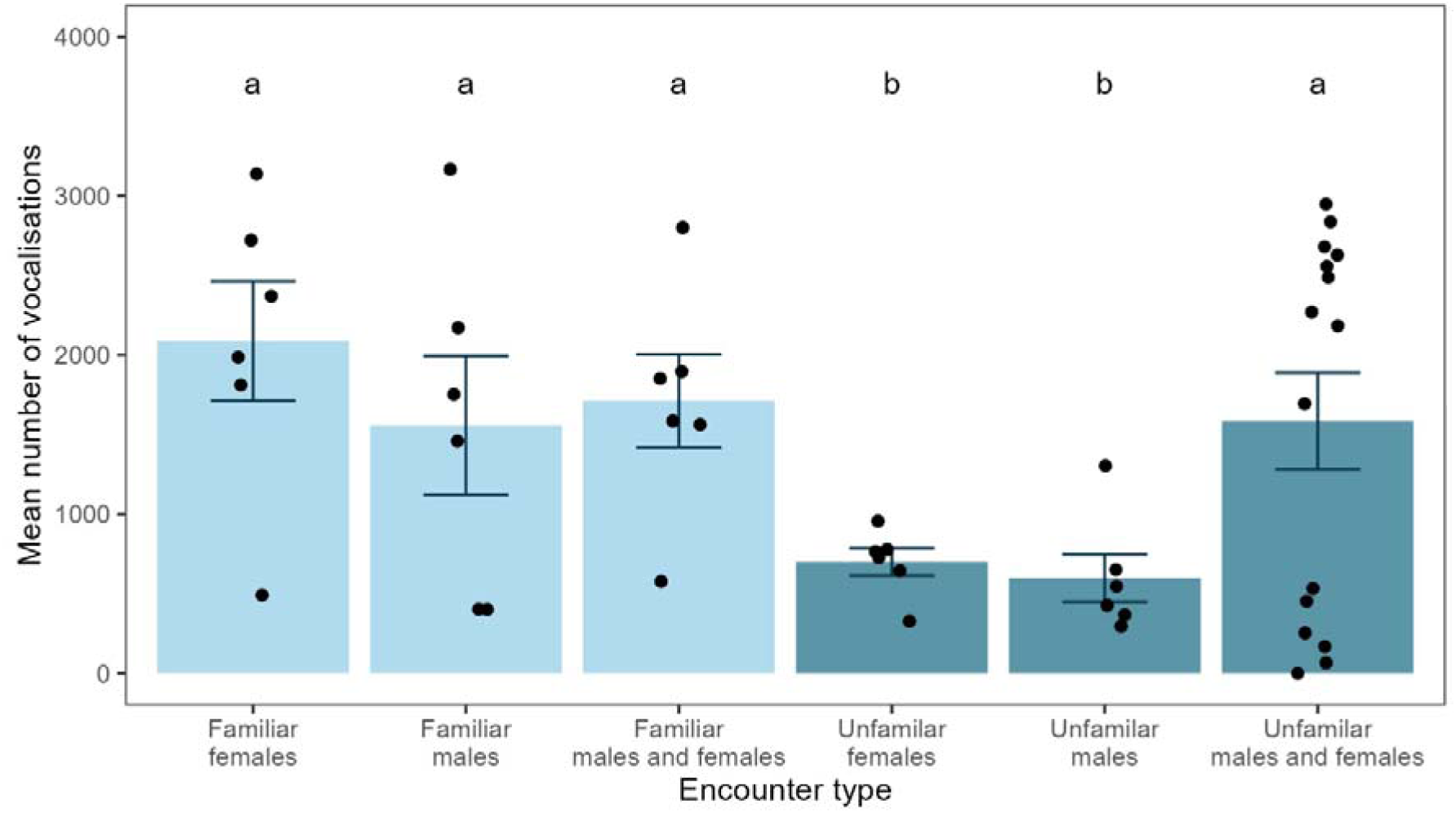
Mice vocal production depends on social context. Each bar chart represents the average number of vocalisations (+/- standard error) emitted by a pair of mice during ten-minute encounters between two individuals, familiar or unfamiliar, of the same or different sex (n = 15 for unfamiliar males and females encounter, and n = 6 for other encounter types). The letters a and b indicate contexts with a significantly different mean number of vocalisations (Bayesian multilevel model: *number of vocalisations ∼ encounter type*, using gaussian family). We recorded on average more vocalisations when the two individuals, female or male, were already living together ("familiar" contexts) than if they had never met before (“Unfamiliar” contexts), unless when the two unfamiliar individuals were of different sexes. In the latter case, the distribution of vocalisations was bimodal: there were either a high number of vocalisations (between 2,000 and 3,000), or a low number (between 0 and 500).

### The African striped mouse vocal repertoire

We detected and categorized recorded vocalisations using Deepsqueak© software. We performed a k-means classification based on the frequency contour and duration of the vocalisations. This unsupervised method separated the vocalisations into seven clusters characterized by particular spectro-temporal parameters (Figure 3). Table 2 summarizes the mean values of these acoustic parameters for each cluster. Most mouse vocalisations were characterized by narrow frequency bands, with varying degrees of frequency modulation. For five of the clusters, frequency modulation was either zero ("flat" cluster), rising ("up" cluster), falling ("down" and "longdown" clusters) or S-shaped ("modulated" cluster). The "trill" and "longdown-trill" clusters were distinguished from the others by a rapid, pronounced sinusoidal frequency modulation, which characterizes all or part of the signal.

**Figure 3.**
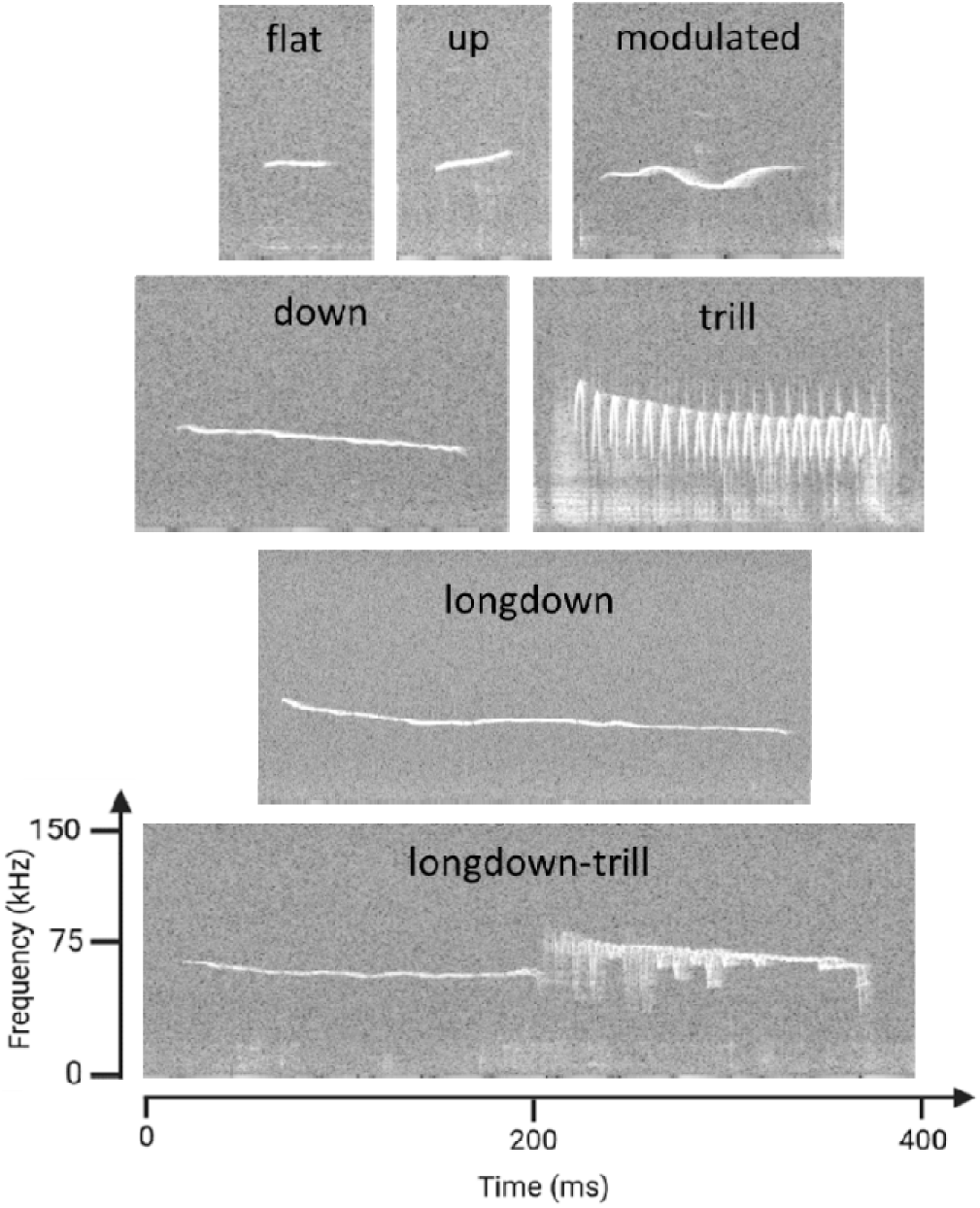
The vocal repertoire of the African striped mouse. Unsupervised K-means clustering on frequency contours of 63,694 recorded vocalisations led to a repertoire of seven acoustic clusters. The acoustic characteristics of these clusters are detailed in Table 2.

**Table 2.** Spectro-temporal features of the African striped mouse vocalisations. Values are the mean of acoustic measurements for each vocalisation type ± standard deviation (extracted from 63,694 vocalisations).

| *Vocalisation type* | *Number of calls* | *Duration (ms)* | *Main Frequency (kHz)* | *Lowest Frequency (kHz)* | *Highest Frequency (kHz)* | *Frequency Bandwidth (kHz)* |
| --- | --- | --- | --- | --- | --- | --- |
| flat | 41,713 | 25.5 ± 1.2 | 59.6 ± 5.1 | 57.9 ± 5.4 | 61.8 ± 5.3 | 3.9 ± 3.0 |
| up | 8,437 | 36.5 ± 1.2 | 59.1 ± 2.5 | 57.3 ± 2.8 | 64.1 ± 2.9 | 6.7 ± 3.0 |
| modulated | 6,455 | 69.8 ± 1.6 | 58.1 ± 3.0 | 55.7 ± 3.2 | 61.1 ± 3.6 | 5.4 ± 3.3 |
| down | 3,174 | 128.5 ± 2.5 | 54.9 ± 4.5 | 51.1 ± 5.2 | 60.5 ± 5.2 | 9.3 ± 4.7 |
| trill | 2,435 | 77.3 ± 4.5 | 74.3 ± 8.4 | 64.8 ± 10.8 | 83.1 ± 8.0 | 18.3 ± 10.4 |
| longdown | 1,203 | 266.1 ± 11.9 | 56.6 ± 6.6 | 50.1 ± 6.7 | 68.1 ± 10.1 | 18.0 ± 10.5 |
| longdown-trill | 277 | 249.9 ± 9.7 | 64.7 ± 7.7 | 50.8 ± 6.9 | 78.0 ± 7.0 | 27.2 ± 8.2 |

### Social context drives the use of vocal repertoire

As illustrated in Figure 4, the vocal repertoire deployed by the mice depended on the sex and familiarity of the individuals in contact. The mice emitted "flat" vocalisations in all contexts (shown in red in Figure 4), and these calls account for most of the vocalisations emitted when two unfamiliar males meet (94.8%). In other contexts, mice used a more varied repertoire. This variation was particularly true during encounters between unfamiliar females and males. In this context, the mice produced 63.1% "flat" calls (other contexts: 71-94.8%), but compared to all other contexts more “down” calls (6.6% vs. 0.1-3.8%), more “longdown” calls (2.7%, vs. 0.0-1.5%), more trills (6.6% vs.0.0-0.2%) and more longdown trills (0.8% vs. 0.0%). The mice also used a diversified repertoire when familiar females and males met, with more “down” calls (3.8%), “longdown” (1.5%) and a few trills (0.2%), than when unfamiliar individuals of the same sex met. Modulated calls were emitted in greater proportion during encounters between individuals of the opposite sex, whether unfamiliar (10.5%) or familiar (7.3%), than during encounters between same-sex individuals. However, the proportion of modulated calls was higher when the individuals are of the same sex but familiar than when they have never met (4.5% for familiar males and 5% for familiar females vs. 0.3% and 1.2% for unfamiliar individuals, respectively for each sex). In encounters between familiar females, there was a significant proportion of “up” calls (22.3%) compared to other encounters (see supplementary figure 1 for all comparisons).

**Figure 4.**
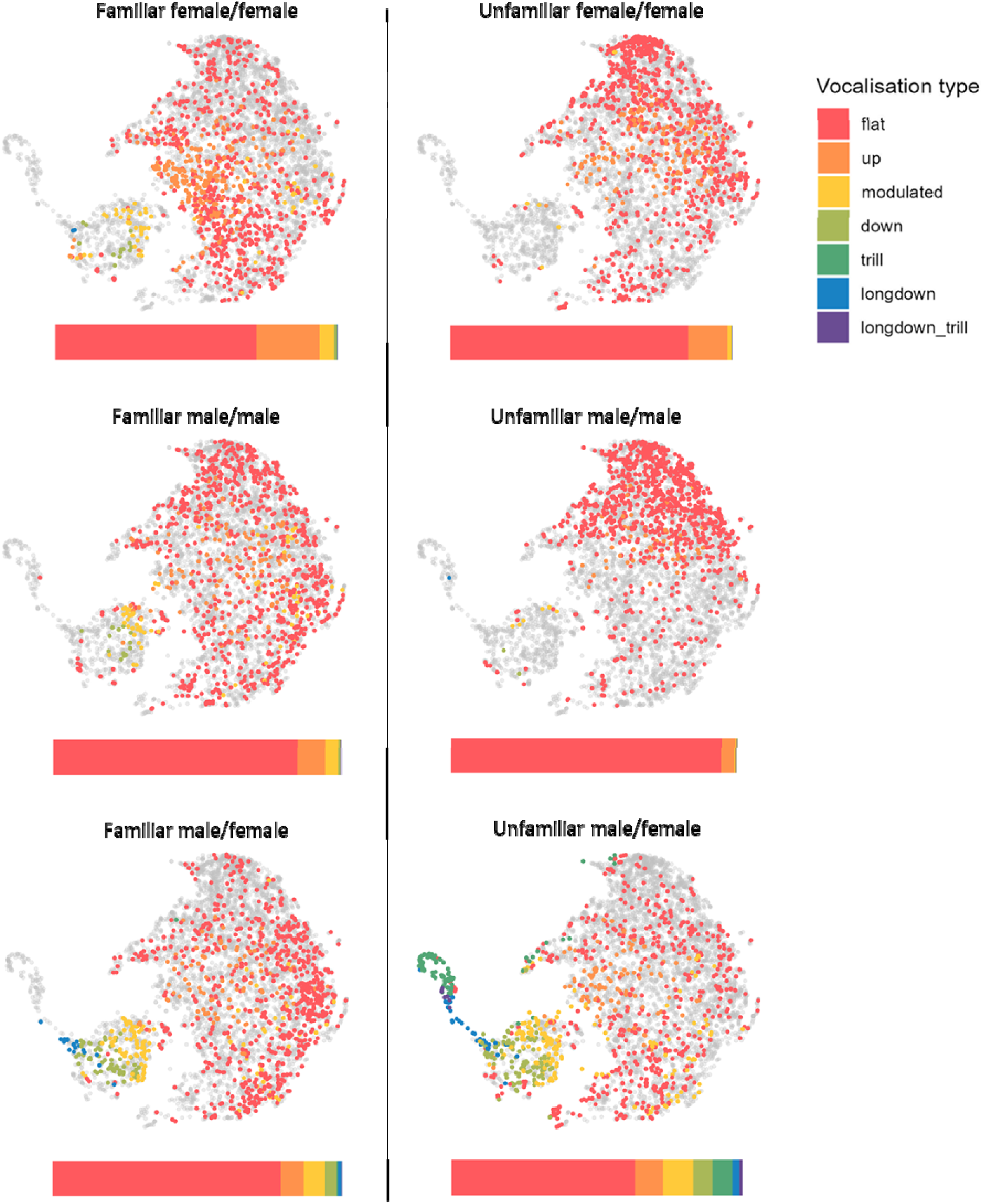
The use of the vocal repertoire by the African striped mouse is context-dependent. Each panel is an acoustic space constructed using UMAP algorithm of dimension reduction applied on 9 acoustic parameters extracted with Deepsqueak software. Each dot represents one vocalisation (for clarity, only 1500 vocalisations randomly selected among all vocalisations detected within an encounter type are represented). The coloured labels correspond to the repertoire illustrated in Figure 3 for calls of each respective encounters, where calls of other encounters are represented by grey dots. Bar charts under the UMAP representation show the distribution of calls for each type of encounter.

### Number of vocalisations and proportion between call types do not change over time

We measured the number of vocalisations and their distribution between the different vocalisation types during the 10 minutes of recording, for each of the six types of encounter (Figure 5). In all contexts, we observed a short adaptation time: during the first minutes, the mice emitted few calls, but their rate of vocal production reached its regular level as early as the third or fourth minute. The rate of vocal production remained remarkably stable over time, except during encounters between same-sex unfamiliar mice, for which we observed a progressive decrease in the number of vocalisations (Bayesian model: percentage of calls per minute ∼ Time * Type of encounter + (1 | Identity of animals); for males, we observed an average decrease of call rate of 1.07% per minute, 95% CI [0.52%; 1.74%], and 1.14% [0.48%; 1.70%] for females. In all social contexts, the relative proportion of different vocalisation types was remarkably stable over time (Figure 5), emphasizing that each context induces a particular use of the mice’s vocal repertoire (Bayesian model: percentage of calls ∼ Time * Type of vocalisation + (1 | Identity).

**Figure 5.**
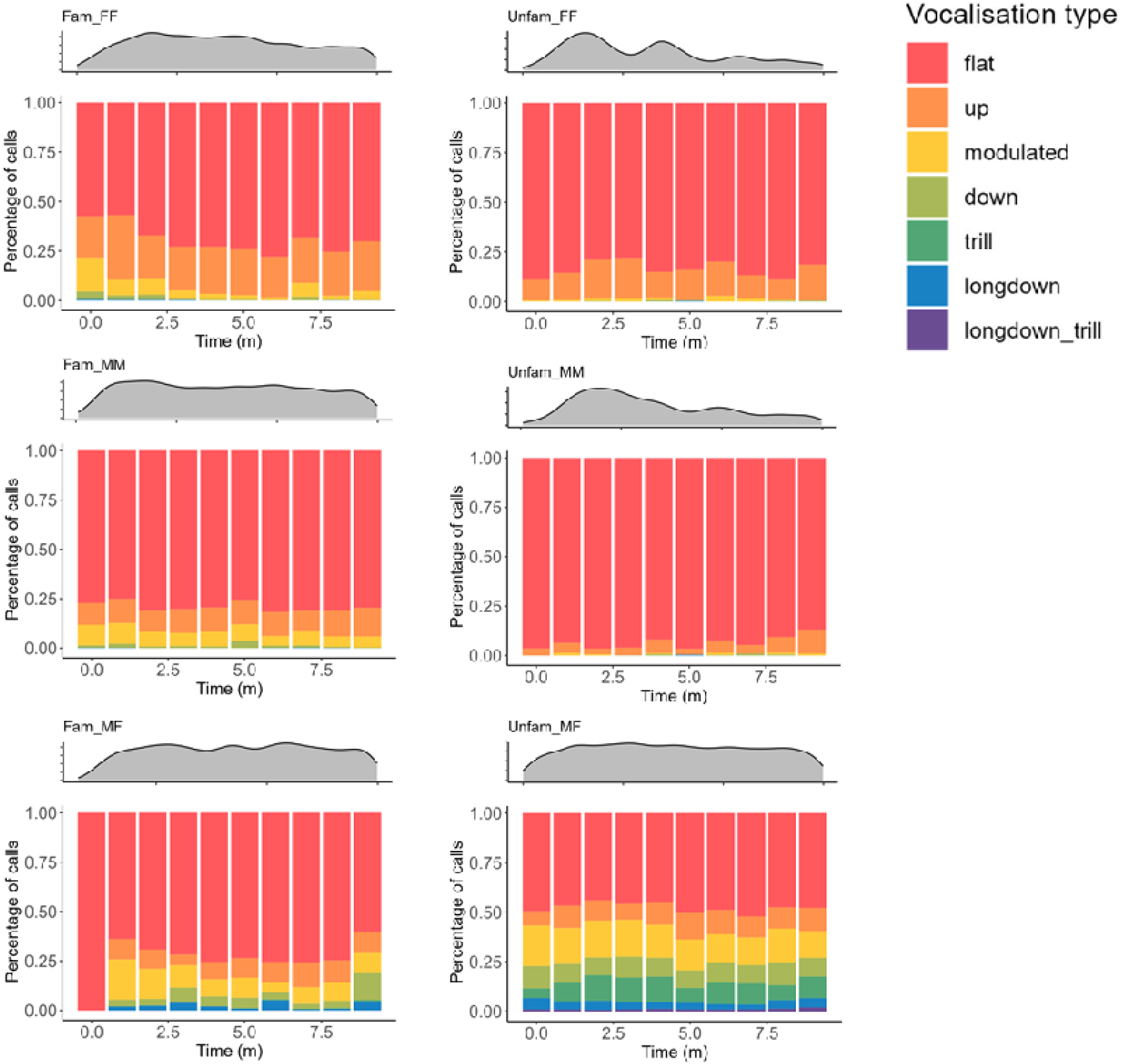
Temporal dynamic of vocalisations as a function of social context. The gray curves represent the evolution of the number of vocalisations per minute over the ten minutes of recording. Colored bar graphs represent the average percentage of each type of vocalisations.

### Familiar males and females display turn-taking

We measured the number and proportion of temporally overlapping vocalisations in each recording context. The results show that overlaps between vocalisations were rarest during encounters between familiar females and males, while they were the most frequent during encounters between unfamiliar females and males (Figure 6; see Table 3 for comparisons of the percentages of overlapping vocalisations between all encounter types; see Figure 1 which shows that the total number of vocalisations was not significantly different between the two conditions; see also supplementary Figure 2 which confirms that the total duration of vocalisations was not significantly different between the two conditions).

**Figure 6.**
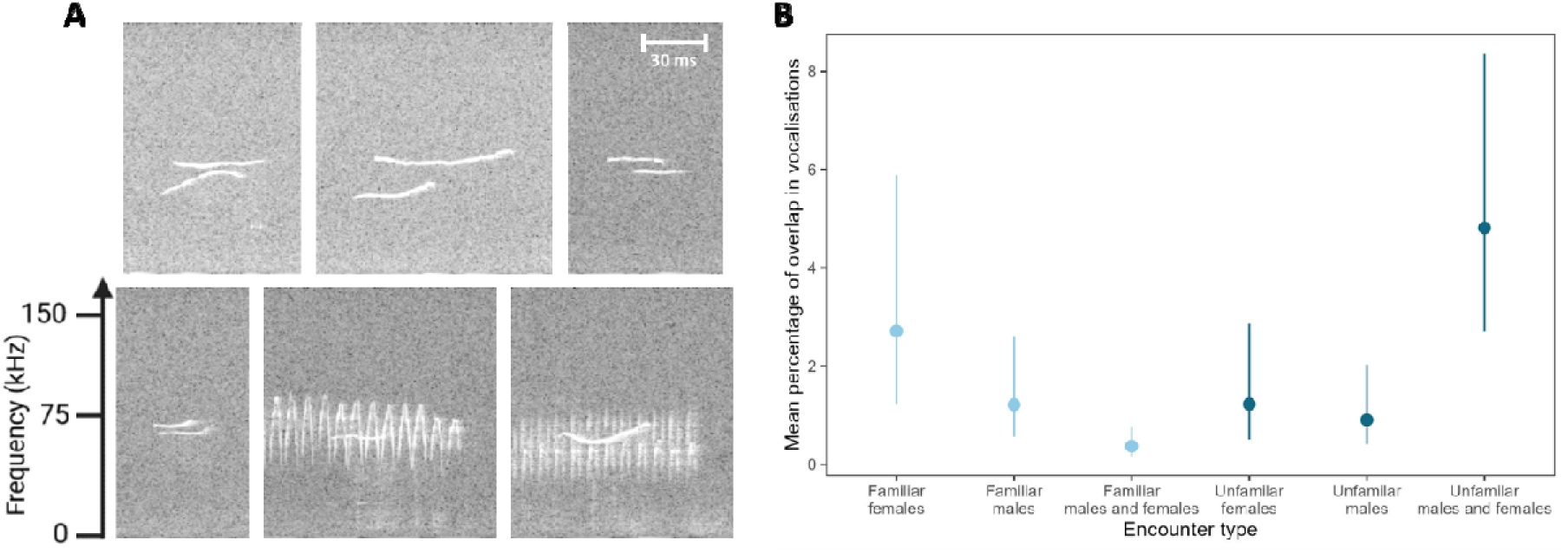
Vocal overlapping during mice encounters. **A.** Examples of overlapping vocalisations. **B.** Proportion of overlapping calls for each encounter type. There is virtually no overlap between calls during encounters between a familiar female and a familiar male. The highest proportion of overlap is observed in encounters between an unfamiliar female and an unfamiliar male. Dots represent the medians of the posterior distributions and bars represent 95% CI of the data fitted using the Bayesian model *Percentage of overlapping calls ∼ Encounter type* (gaussian distribution).

**Table 3.** Less vocal overlapping is found when mice are familiar. The table presents contrasts of the encounter type in rows compare to the encounter type in column, with their 95% credible interval.

|  | Familiar females | Familiar males | Familiar males and females | Unfamiliar females | Unfamiliar males | Unfamiliar males and females |
| --- | --- | --- | --- | --- | --- | --- |
| Familiar females |  | 1.4%, [-0.6%, 4.7%] | 2.1%, [0.8%, 5.5%] | 1.3%, [-0.7%, 4.7%] | 1.6%, [0%, 4.9%] | -2%, [-5.9%, 1.7%] |
| Familiar males | -1.4%, [-4.7%, 0.6%] |  | 0.7%, [0%, 2.4%] | 0%, [-1.9%, 1.7%] | 0.2%, [-1%, 1.9%] | -3.4%, [-7.3%, -1.1%] |
| Familiar males and females | -2.1%, [-5.5%, -0.8%] | -0.7%, [-2.4%, 0%] |  | -0.7%, [-2.6%, 0%] | -0.5%, [-1.7%, 0.1%] | -4.1%, [-8%, -2.3%] |
| Unfamiliar females | -1.3%, [-4.7%, 0.7%] | 0%, [-1.7%, 1.9%] | 0.7%, [0%, 2.6%] |  | 0.2%, [-1%, 2.1%] | -3.4%, [-7.3%, -0.8%] |
| Unfamiliar males | -1.6%, [-4.9%, 0%] | -0.2%, [-1.9%, 1%] | 0.5%, [-0.1%, 1.7%] | -0.2%, [-2.1%, 1%] |  | -3.6%, [-7.6%, -1.5%] |
| Unfamiliar males and females | 2%, [-1.7%, 5.9%] | 3.4%, [1.1%, 7.3%] | 4.1%, [2.3%, 8%] | 3.4%, [0.8%, 7.3%] | 3.6%, [1.5%, 7.6%] |  |

## Discussion

Here we reveal how a mouse uses its diverse vocal repertoire according to social context. We found that the striped mouse has a diverse vocal repertoire, which it masters both quantitatively and qualitatively. We showed that the content of vocal exchanges between two striped mice differs according to whether the two mice are individuals who already know each other (sisters, brothers or a paired female and male), or who have never seen each other (two females, two males or unfamiliar female and male). In short, our results provide a fresh perspective on the richness of rodent vocal exchanges, compared with that provided by laboratory mice and rats, where studies are usually focused on limited, unnatural interactions, restraining the way animals use their vocal repertoire.

African striped mice vocalized least during encounters between unfamiliar individuals of the same sex. In this context, their vocal production was essentially limited to flat calls. The mice appear to establish vocal contact during the first few minutes of the encounter, with a marked increase in the number of vocalisations emitted per minute, before rapidly decreasing their vocal production. These results converge with observations made in the field: a coincidental encounter between two African striped mice from different groups results in agonistic interactions, after which the animals rapidly move away from the conflict zone (51,57). Our results are congruent with previous studies on female house mice *Mus musculus musculus*, where two unfamiliar females vocalized less during the first night of contact compared to the second night (32). For male house mice, it has also been shown that urine from unfamiliar males elicit less vocalisations than urine from familiar males (31). Therefore, we might consider that only a few minutes of contact, including a short vocal exchange, is enough to resolve the interaction between unfamiliar individuals of the same sex.

The situation became different if the two unfamiliar striped mice that are put together were of different sexes. Here we observed a bimodal pattern: either the number of vocalisations was very low, or it was very high. In the latter case, the rate of vocal production remained sustained throughout the encounter, and the vocalisations were diverse, using the whole vocal repertoire, particularly modulated vocalisations and trills. This bimodality could be explained by differences in the sexual motivation of the individuals. It has been shown that male house mice produce more vocalisations in response to the presentation of mature, reproductive females than when the females are not receptive (33). It is likely that some of our female striped mice were sexually receptive (oestrus) at the time of the experiments while others were not. The oestrus cycle of striped mice is relatively long with 11 days, 5 of which are in oestrus (58), which could well explain our bimodal distribution, as by chance we would expect about 45% of the females to be in oestrus. It is also possible that individuals brought into contact may or may not be attracted to each other. In laboratory breeding facilities, we have observed that some pairs of mice reproduce without difficulty, while others seem to ignore each other and do not reproduce (this latter situation concerns more than one pair out of two).

When two familiar individuals meet after having been separated, our experiments show that their vocal production increases rapidly and remains high throughout the encounter. This continuity in vocal production is in line with what is observed in the wild African striped mice, when the individuals emerge from the nest at first sunlight and interact for around 30 minutes (51,52). Our results show that during encounters between familiar individuals, their vocal repertoire is diverse and does not differ according to whether the two mice are two females or two males. The only minor difference was the “up” vocalisation that was more present during encounters between familiar females. We found no difference in repertoire complexity between females and males. In this context, the richest vocal repertoire was observed during encounters between familiar females and males, with a high production of modulated calls, up, down and longdown. Trills, on the other hand, were almost virtually absent, marking a clear difference in the vocal repertoire used compared to encounters between unfamiliar females and males, where the repertoire used was the most diverse. Remarkably, this repertoire (the relative quantities of each type of vocalisation) remained stable throughout the 10-minute experiment.

Overall, these results demonstrate that African striped mice are capable of identifying the sex and familiar/unknown identity of their conspecifics. Behavioural studies already indicate that striped mice are capable of individual recognition, preferring specific group members over others, and showing an increased semipositive reaction when their preferred partner is removed for one day and then returned (52). This identification appears to be rapid, occurring within the first few minutes of the experiment, before the individuals have had time to interact vocally in any meaningful way. It is generally assumed that the identification of sex and status occurs primarily via the olfactory channel (59). Identification of sex by odour is thus strongly suggested by the female-male unfamiliar trials: despite the fact that these mice have never met in the past, they seem to recognize that they are of opposite sex at the very beginning of the trial. Nonetheless, we cannot rule out an additional, rapid identification of a precise sexual and individual vocal signature that could be encoded in the calls.

Understanding the biological significance of these calls will require further studies where individual behaviours are documented in parallel with the vocalisations produced, ideally by precisely identifying individual vocal contributions during interactions. However, the present work already brings some interesting insights. Firstly, it should be noted that the vocalisations defining the vocal repertoire actually line up along an acoustic continuum, with the exception of the trill, which has acoustic features that distinguish it from other vocalisations. Most of the vocalisations of the African striped mouse therefore constitute a graded vocal repertoire. This acoustic gradation could encode information of a "dynamic" type, i.e. linked to short-term fluctuations in the sender’s physiological and psychological states (2,60,61). Graded vocal signals are common in animals, particularly in species whose individuals live in social groups and exchange subtle dynamic information (62). For example, the five most common tonal vocalisations of the bonobo *Pan paniscus* (high-hoot, bark, soft bark, peep-yelp, and peep calls) form an acoustic continuum that reflects the arousal level of the sender (2). In our striped mouse, the up, modulated and down calls define an acoustic continuum around the flat vocalisation. This continuum could enable mice to encode dynamic information (level of arousal and/or valence). This hypothesis is supported by the fact that these vocalisations are used differently depending on the social context. Thus, we can assume that the use of up, modulated and down vocalisations in female-male encounter contexts reflects a high level of arousal on the part of the senders. As mentioned above, these modulated calls may enable animals to produce more complex, less redundant patterns, which could increase the likelihood that these calls contain more information (cf. 52, on the Mathematical Theory of Communication). This hypothesis is reinforced by the fact that it is in these female-male contexts - particularly when the individuals are unfamiliar - that the most trills are recorded.

Our results clearly show that the African striped mouse adjusts its vocal repertoire according to the social context in which it is immersed. We also found that these mice adjust the timing of their vocalisations based on whom they are interacting with. The mice generally do not overlap their calls with a neighbour’s, but the incidence of overlapping declines from a standard 5% in male/female pairs of previously unfamiliar individuals to less than 1% in familiarized male/female pairs. Overlap avoidance may be a means by which animals facilitate exchange of information (64), whereas a tendency to overlap could signify aggressiveness (65) – which might be relatively higher between unfamiliar mice. Control over interactive timing suggests that ‘intention’ is present in some vocalisations (66,67) see also (68).

One critical limitation in our study, and those of laboratory mice (*M. musculus*) until now, is that we could not identify which individual of the pair vocalized. This constraint arises because the mice – *R. pumilio*, and *M. musculus* as well – do not give any visible indication when vocalizing, and the very short distance between paired individuals does not permit convenient application of a triangulation method, using multiple microphones, for precisely localizing the source of each vocalisation in an encounter trial. Thus, all of our data reflect behaviour of dyads as opposed to specific individuals. But recent technical developments (69,70) may allow us to overcome this drawback in future studies of *R. pumilio*. Another recent development with this species is the assembly of a high-quality reference genome (71) of this wild mouse species. It is expected that the striped mice genome will eventually facilitate deciphering the genetic architecture of the complex vocal communication we are beginning to report.

## Acknowledgements

This study has been funded by the University of Saint-Etienne, the Institut universitaire de France, the CNRS, the Inserm and the Labex CeLyA.

---

**Supplementary Figure 1.**
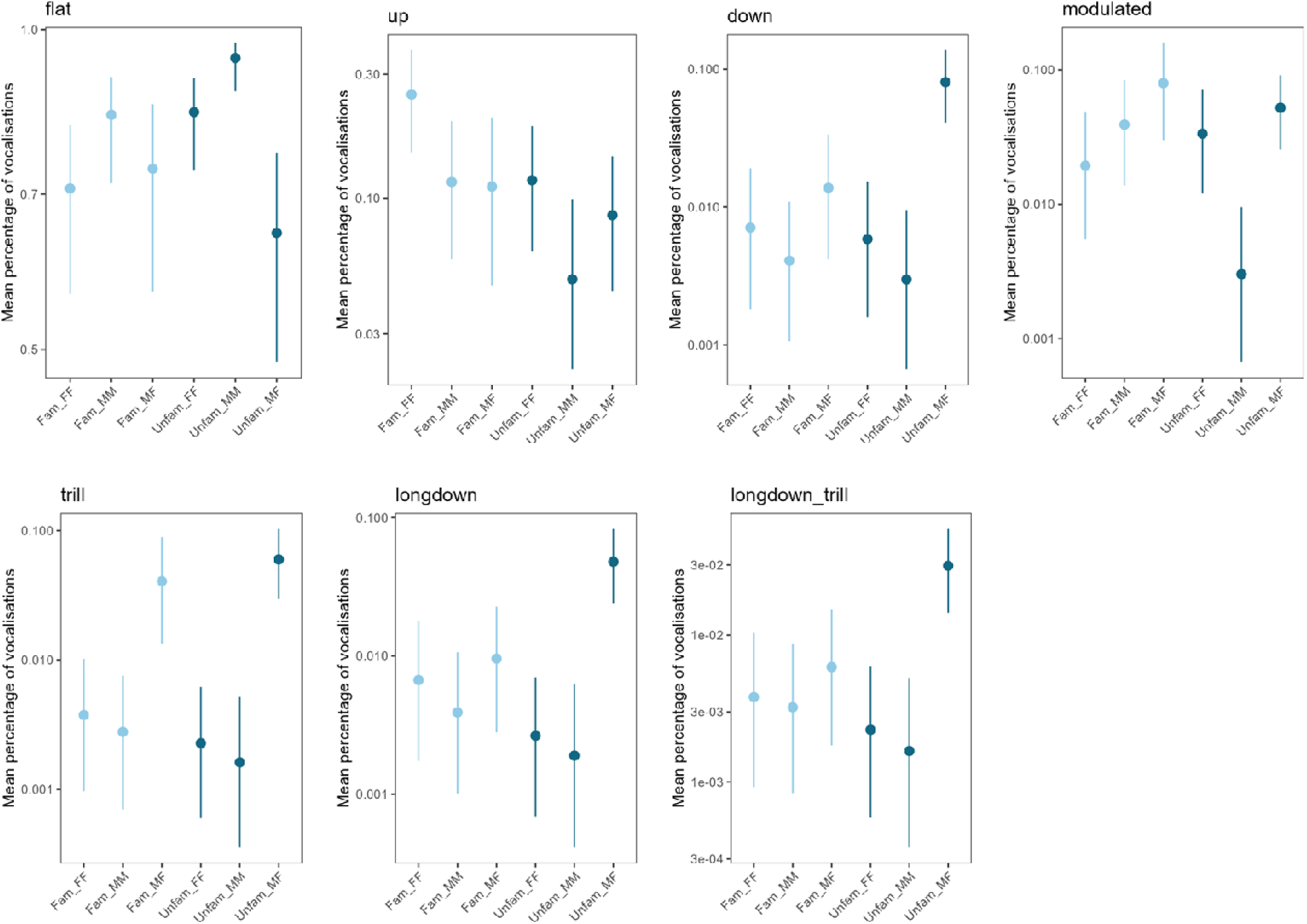
The relative proportion of vocalisation types depends on social context. Fam_FF: familiar females; Fam_MM: familiar males; Unfam_FF: unfamiliar females; Unfam_MM: unfamiliar males; Unfam_MF: unfamiliar female and male. Solid disks represent the medians of the posterior distributions and bars represent 95% CI of the data fitted using the Bayesian model *Percentage of calls ∼ Encounter type*. Model fitted using a Dirichlet distribution.

**Supplementary Figure 2.**
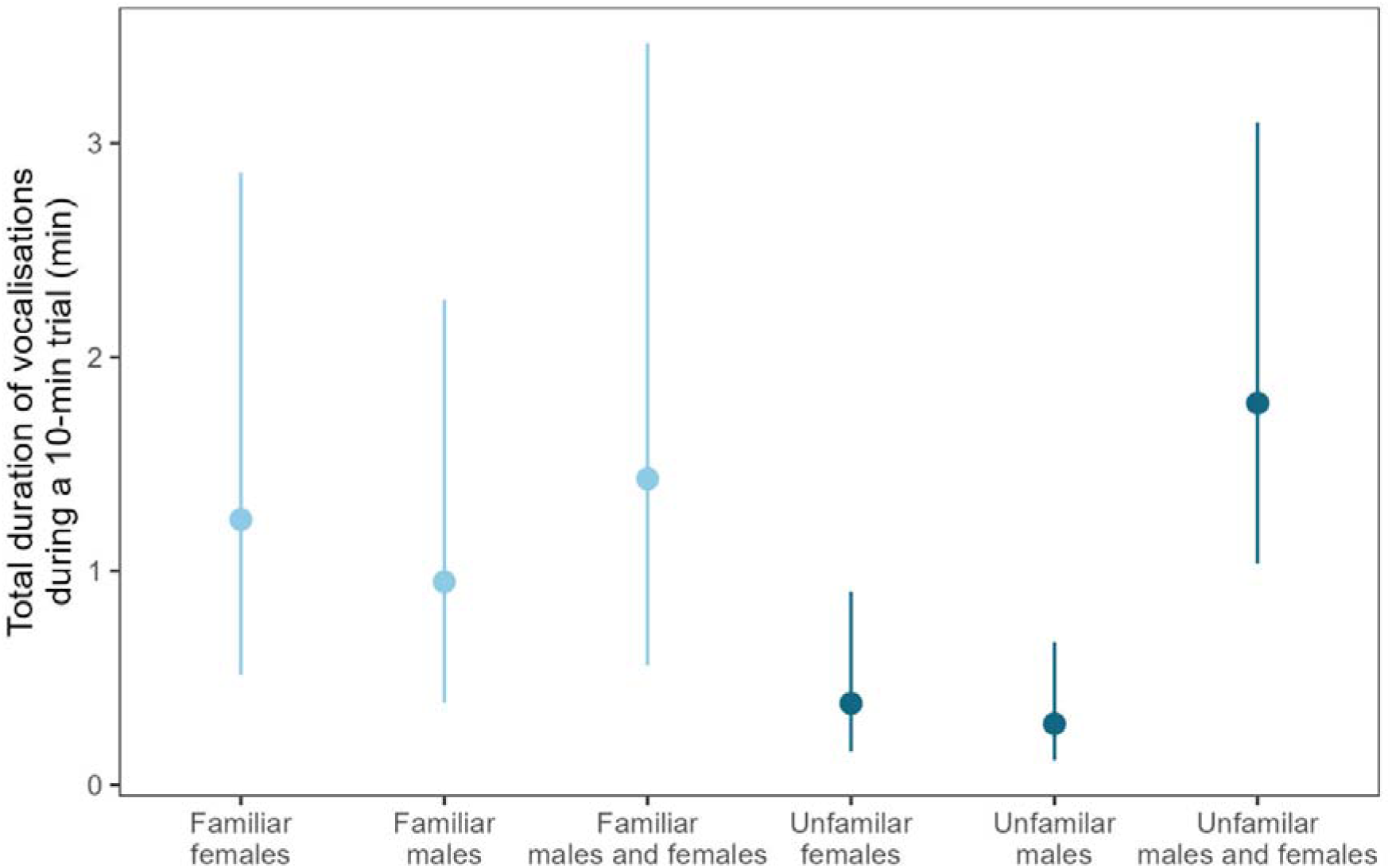
Total duration of vocalizations depending on social context. Solid disks represent the medians of the posterior distributions and bars represent 95% CI of the data fitted using the Bayesian model *Total duration of vocalizations over a 10-min trial ∼ Encounter type*. Model fitted using a gamma distribution.

