## Supplementary Material for "A cooperatively breeding mouse shows flexible use of its vocal repertoire according to social context"

**Supplementary figure**

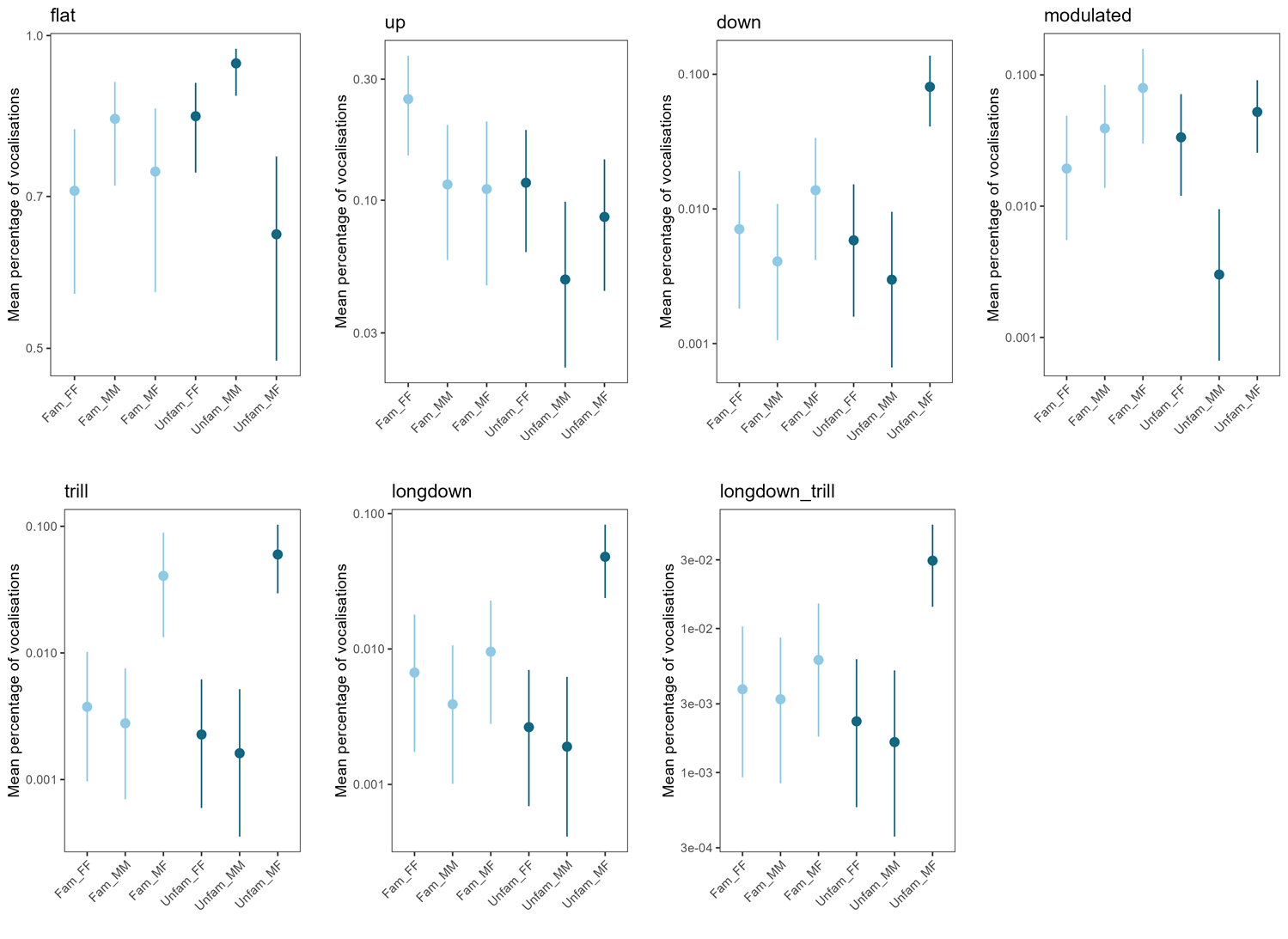

**Supplementary Figure 1. The relative proportion of vocalisation types depends on social context.** Fam_FF**:** familiar females; Fam_MM: familiar males; Unfam_FF: unfamiliar females; Unfam_MM: unfamiliar males; Unfam_MF: unfamiliar female and male. Solid disks represent the medians of the posterior distributions and bars represent 95% CI of the data fitted using the Bayesian model *Percentage of calls ~ Encounter type*. Model fitted using a Dirichlet distribution.

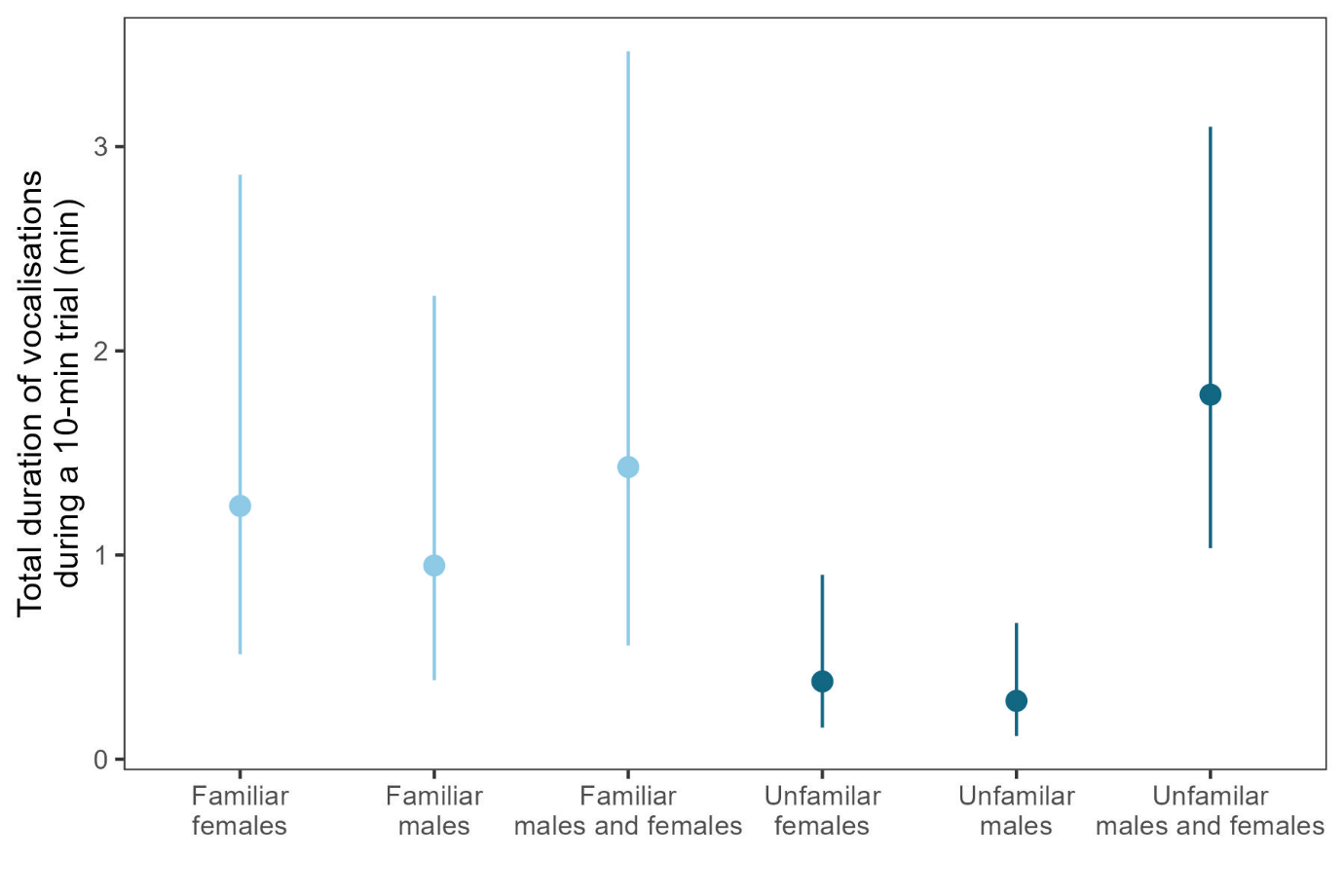

**Supplementary Figure 2. Total duration of vocalizations depending on social context.** Solid disks represent the medians of the posterior distributions and bars represent 95% CI of the data fitted using the Bayesian model *Total duration of vocalizations over a 10-min trial ~ Encounter type*. Model fitted using a gamma distribution.

**Data availability**

For double-blind review reason, data are currently available under an anonymous repository: <https://zenodo.org/records/11083441>. In case of publication, all data will be available on the author’s GitHub.
